# Spatiotemporal EP4-fibulin-1 expression is associated with vascular intimal hyperplasia

**DOI:** 10.1101/2023.11.09.566500

**Authors:** Shigekuni Okumura, Sayuki Oka, Takako Sasaki, Marion A. Cooley, Yuko Hidaka, Shota Tanifuji, Mari Kaneko, Takaya Abe, Richard M. Breyer, Hiroshi Homma, Yuko Kato, Utako Yokoyama

## Abstract

**Aims:** Cyclooxygenase-2– and microsomal prostaglandin E synthase-1–derived prostaglandin E_2_ (PGE_2_) are involved in vascular intimal hyperplasia (IH). Although extensive studies have revealed the roles of PGE_2_ receptors (EPs) in IH, spatiotemporal EP expressions and downstream targets have not been fully elucidated. In this study, we focused on EP4 and investigated its role in vascular IH.

**Methods and Results:** We generated EP4 reporter mice (*Ptger4*-IRES-nlsLacZ) and found prominent EP4 expression in the proliferative neointima 2 weeks after femoral artery wire injury. Expression of EP4 were returned to the baseline level 4 weeks after vascular injury (VI). Injury-induced IH was diminished in vascular smooth muscle cell (VSMC)-specific EP4 heterozygous deficient mice (*Ptger4*^fl/+^;*SM22*-*Cre*) 2 and 4 weeks after VI compared to *SM22*-*Cre*, whereas injury-induced IH was exacerbated in VSMC-specific EP4-overexpressing mice (*Ptger4*-Tg) compared to controls (non-Tg). Systemic EP4 antagonist administration reduced VI-induced IH in wild-type mice. We investigated the role of extracellular matrix proteins, as downstream regulated targets of EP4. Stimulation of EP4 increased mRNA and protein levels of fibulin-1 (a multifunctional glycoprotein) in *Ptger4*-Tg VSMCs. Fibulin-1C or -1D recombinant proteins increased VSMC proliferation, whereas proliferation was decreased in fibulin-1–deficient VSMCs. We generated multiple deletion mutants of fibulin-1C and found that EGF-like modules 6-8 appear to be involved in fibulin-1–mediated proliferation. Among binding partners of fibulin-1, extracellular matrix protein 1 (ECM1) was upregulated by EP4 stimulation, and fibulin-1 and ECM1 proteins additively enhanced VSMC proliferation. Similar to EP4 expression, both fibulin-1 and ECM1 were abundantly expressed in the neointima 2 weeks after VI. Furthermore, injury-induced IH was attenuated in VSMC-specific fibulin-1 deletion mice (*Fbln1*^fl/fl^;*SM22*-Cre) compared to *Fbln1*^fl/fl^.

**Conclusions:** EP4 was upregulated in proliferative IH, and EP4-induced fibulin-1 cooperated with ECM1 to promote IH through VSMC proliferation. The calcium binding EGF-like modules 6-8 of fibulin-1 are indicated to regulate cell proliferation.

**A Translational Perspective:** Recent advances in drug-eluting stents have significantly contributed to the reduction of vascular IH. However, the detailed mechanism underlying IH after stenting remains to be elucidated. We found that prostaglandin E_2_-EP4–induced fibulin-1 plays a role in IH through VSMC proliferation. It is well recognized that prostaglandin E_2_ plays a role in IH, but inhibition of cyclooxygenase-2 has side effects such as thrombogenesis. Because EP4 and fibulin-1 were upregulated specifically in the neointima after vascular injury, oral or local administration of an EP4 antagonist or the downregulation of fibulin-1 would be potential therapeutic strategies to restrain IH.

## 1. Introduction

Vascular restenosis is a wound-healing response to mechanical injuries, such as balloon angioplasty and stenting. A large-cohort study demonstrated that restenosis 6 to 8 months after coronary stenting is correlated with 4-year mortality. ^1^ Once the blood vessels are injured, disruption of the endothelial tight junction allows inflammatory cytokines, chemokines, and several growth factors to promote vascular smooth muscle cell (VSMC) proliferation and migration from the tunica media toward the internal lumen. ^2^ The intimal hyperplastic area then undergoes repeated synthesis and degradation of extracellular matrices (ECM), eventually resulting in a high percentage of collagen- and proteoglycan-rich ECM. ^2^

Compared to standard balloon angioplasty, drug-eluting stents (DES) have significantly reduced restenosis rates. A DES is a metal stent coated with a drug that inhibits VSMC proliferation. First-generation DES used sirolimus, paclitaxel, and other drugs that inhibit cell growth and proliferation; second-generation DES use a derivative of sirolimus as the carrier drug, allowing for lower drug concentration and lesser toxicity. ^3^ However, despite advances in stent technology, cardiovascular adverse events, including restenosis, still occur in approximately 5% to 10% of all patients. ^4^

Prostaglandin E_2_ (PGE_2_) is synthesized by cyclooxygenase-2 (COX-2) in the vessels after vascular injury. ^5^ Activation of the PGE_2_-producing enzymes COX-2 or microsomal PGE_2_ synthase-1 (mPGES-1) exacerbates injury-induced vascular intimal hyperplasia (IH). ^5, 6^ However, clinical studies have revealed that using COX-2 inhibitors to inhibit PGE_2_ production is not suitable for long-term use because of side effects such as thromboembolism and gastric ulcers. ^7^ There are four types of PGE_2_ receptors (EP1-4), and their roles have been extensively studied using genetically modified mice and receptor selective compounds. It has been demonstrated that EP2 inhibited injury-induced IH via attenuation of VSMC proliferation and migration, ^8^ while EP3 promoted directional VSMC migration and IH, and EP4 in VSMCs contributed to enhancement of IH. ^9^ In addition, EP4 signaling in endothelial cells reportedly accelerated endothelialization of injured vessels and restrained IH. ^10^ Hence, the roles of EP4 in IH are considered to depend on the cell type. These data suggest that a variety of mechanisms, depending on receptor and cell type, underlie vascular IH. However, spatiotemporal EP expression and roles in IH remain to be elucidated.

ECMs are essential for regulating VSMC proliferation and migration, which are the key molecular mechanisms in IH. ^11^ Glycoproteins such as laminin, nidogen, fibronectin, and tenascin bind to other molecules, such as collagen, to stimulate VSMC proliferation and migration. Proteoglycans such as versican and hyaluronan form insoluble complexes that promote VSMC proliferation. ^11^ Tenascin-C has been implicated to play a role in VSMC proliferation and migration in vascular IH. ^6, 9^ However, a study using mPGES-1–deficient mice raises the possibility that other ECMs, besides tenascin-C, contribute to VSMC-mediated IH. ^6^ It has also been demonstrated that EP4 promotes physiological IH during development via ECM production, including the glycoprotein fibulin-1 and hyaluronic acid. ^12, 13^ Based on these findings, we focused in this study on the roles of EP4 to promote changes in the ECM in injury-induced pathological vascular IH.

## 2. Materials and methods

### 2.1 Reagents

An EP4 agonist (ONO-AE1-437) was kindly provided by Ono Pharmaceutical Company (Osaka, Japan). An EP4 antagonists (CJ42794, ONO-AE3-208) were purchased from Cayman Chemicals (Ann Arbor, MI, USA). Prostaglandin E_2_ (PGE_2_) and indomethacin were obtained from Calbiochem (Billerica, MA, USA) and Tokyo Chemical Industry (Tokyo, Japan), respectively.

An antibody for fibulin-1 (NBP1-84725) was purchased from Novus Biologicals (Centennial, CO, USA). An antibody for ECM-1 (11521-1-AP) was purchased from Proteintech Group Inc. (Chicago, IL, USA). An antibody for versican GAGβ (AB1033) was obtained from Sigma-Aldrich (St. Louis, MO, USA). Antibodies for PCNA (ab92552) and α-smooth muscle actin (ab5694) were obtained from Abcam (Cambridge, UK). An antibody for CD68 (MCA1957) was obtained from Bio-Rad (Hercules, CA, USA). An antibody for von Willebrand factor was obtained from Dako (Glostrup, Denmark). Antibodies for rabbit IgG, Alexa Fluor 546 and 647 (A10040 and A21244), and Hoechst 33258 (H3549) were purchased from Invitrogen (Carlsbad, CA, USA).

### 2.2 Animals

The Ethical Committee of Animal Experiments approved the experiments at Tokyo Medical University School of Medicine (R5-084) and the Institutional Animal Care and Use Committee of RIKEN Kobe Branch (A2001-03). Genetically modified *SM22-Cre* mice (Stock Tg [*Tagln-cre*] 1Her/J) were purchased from the Jackson Laboratory (Bar Harbor, ME, USA). Generations of vascular smooth muscle–specific EP4-overexpressing mice (*Ptger4*-Tg) and non-transgenic mice (non-Tg) were as described previously. ^14^ Generation of *Ptger4* flox mice *(Ptger4*^fl/fl^) was as described previously. ^15^ We generated vascular smooth muscle–specific EP4 heterozygous deficient mice (*Ptger4*^fl/+^;*SM22-Cre*) and control mice (*Ptger4*^+/+^;*SM22-Cre*) by crossing *SM22-Cre* mice with *Ptger4*^fl/fl^ mice, as described previously. ^14^

EP4 reporter mice (*Ptger4*-IRES-nlsLacZ) (Accession number: CDB1380K: https://large.riken.jp/distribution/mutant-list.html) were generated using embryonic stem (ES) cells derived from C57BL/6N strain. ^16^ This method involved the creation of embryonic stem (ES) cells containing an internal ribosomal entry site nuclear localization signal LacZ bovine growth hormone polyadenylation (IRES nlsLacZ bGHpA) construct, as well as a neomycin resistance gene surrounded by a loxP sequence upstream of exon 3 in the EP4 expression site (Supplemental Figure 1A). Targeted ES cell clones were then injected into 8-cell stage ICR embryos, and the resulting chimeric mice were crossed with C57BL/6N mice to achieve germline transmission.

Deletion of all variants of fibulin-1 in mice results in perinatal lethality. ^17^ To investigate the roles of fibulin-1 in VSMCs, we generated fibulin-1 flox mice (*Fbln1*^fl/fl^). The targeting vector for the generation of a conditional *Fbln1* deficient mouse was designed by the Knockout Mouse Project (KOMP repository; https://www.komp.org). The designed vector contained a 4.2-kb region of *Fbln1* genomic DNA on the 5′ homologous arm and a 5.5-kb region on 3′ homologous arm (Supplemental Figure 1B). *Fbln1*-targeted ES cells were isolated by KOMP and provided to the Medical University of South Carolina transgenic core facility for injection into C57BL/6 blastocysts. A germline male chimera was obtained and bred to C57BL/6 mice. The resulting F_1_ mice were crossed with an FLPeR deleter strain (B6.129S4-Gt(ROSA)26Sortm1(FLP1)Dym/RainJ; Jackson Laboratory) ^18^ to excise the β-galactosidase (lacZ gene) and neomycin resistance (neo gene) cassettes flanked by FRT sites (Supplemental Figure 1B). Mice carrying the *Fbln1* floxed allele (*Fbln1*^fl/fl^) were crossed with *SM22-Cre* mice to generate vascular smooth muscle–specific fibulin-1 deletion mice (*Fbln1*^fl/fl^;*SM22-Cre*) (Supplemental Figures 1C-D).

### 2.3 Femoral artery injury model

Male mice aged 12 to 24 weeks and female mice aged 15 to 24 weeks were used for the experiments. The surgical procedure was performed using a dissecting microscope model SMZ-800 (Nikon, Tokyo, Japan) under isoflurane inhalation (1.5% at a flow rate of 1L/min) for anesthesia. Transluminal mechanical injury of the femoral artery was induced by inserting a large wire (0.38 mm diameter, C-SF-15-15, Cook, Bloomington, IN, USA), as previously described. ^19^ Euthanasia of the mice was carried out by cervical dislocation after deep isoflurane inhalation (3% at a flow rate of 1 L/min) at a predetermined number of weeks following vascular injury. Afterwards, a 24-gauge intravenous catheter (Terumo, Tokyo, Japan) was inserted through the descending aorta, and a backward flow of phosphate buffered saline (PBS) followed by 10% formaldehyde (Sigma-Aldrich, St. Louis, MO, USA) was introduced using a micro syringe pump (MSD-1D, AS ONE, Osaka, Japan) to maintain a constant reflux pressure for fixation.

### 2.4 X-gal staining

After cervical dislocation of mice with deep isoflurane inhalation (3% at a flow rate of 1 L/min) with vascular injury, a 24-gauge intravenous catheter was inserted through the descending aorta, and a backward flow of PBS was performed. Subsequently, each sample was incubated in a fixative solution containing 1% paraformaldehyde (Wako, Osaka, Japan), 0.2% glutaraldehyde (Wako), and 0.02% NP40 (Wako) dissolved in PBS at 4°C for 1 h. The samples were then subjected to three 5-min washes at room temperature with a solution of 0.02% NP40 dissolved in PBS. Subsequent staining was performed by immersing each sample in a solution containing 4 mM potassium ferricyanide (K_3_Fe(CN)_6_; Wako), 4 mM potassium ferrocyanide (K_4_Fe(CN)_6_; Nacalai Tesque, Kyoto, Japan), 1 M MgCl_2_ (Wako), and 0.1% 4-chloro-5-bromo-3-indolyl-D-galactopyranoside (X-gal, 40 mg/mL in DMSO; Wako) at 37°C for 48 h with shaking. Following staining, each sample was subjected to four 5-min washes with a solution of 0.02% NP40 containing 2 mM EDTA (Nacalai Tesque) dissolved in PBS at room temperature. The samples were then fixed in 10% formalin overnight at 4°C and dehydrated in 70% ethanol at 4°C. Nuclear Fast Red stain was applied for 1 min to visualize the nuclei, followed by a 10-min water wash.

### 2.5 Construction of expression vectors and purification of fibulin-1 recombinant proteins

To facilitate purification, several deletion mutant proteins including full-length fibulin-1C were expressed with His-myc tags at the C-terminus. Full-length fibulin-1C cDNAs without a stop codon were inserted into pBluescript through KpnI and NotI sites, and this plasmid was used to prepare deletion mutant constructs. Final cDNA was cloned into a modified pCEP-Pu expression vector, which contained His-myc tags before the stop codon through KpnI and NotI sites. The cDNAs of ΔEG2-3, ΔEG4-5, and ΔEG2-5 were prepared by partial EcoRV digestion of the full-length fibulin-1C construct as described previously. ^20^ The following oligonucleotides were used as PCR primers: [1: GTCA<u>GATATc</u>GATGAGTGTGCC (EcoRV site is underlined and a lowercase letter shows an introduced mutation); 2, GTCA<u>GAtATc</u>GATGATGAGTGTGTGAC (the lowercase letters show the mutations); 3, GTCAGCGGCCGCGAGCTCTGC; 4, GTCAGGTACCGCCCCATGGAGCGCC; 5, GTCAGCGGCCGCATCTTCGCAGGTGACCC; 6, GTCAGGATCCTTCCGTTGCAGAC; 7, CACTCATCAATATCCACGCAGGTCCTGC; 8, GCAGGACCTGCGTGGATATTGATGAGTG; 9, GTCAGCTAGCCTGTAGCCAGATG; 10, CTCATGGCAAGGCAATTCGCAGGTGACCCC; 11, GGGGTCACCTGCGAATTGCCTTGCCATGAG] For ΔEG2-7 and ΔEG2-8 constructs, an EcoRV site was introduced to the 5′ end of PCR fragments encoding EG8-CT (primers 1 and 3) and EG9-CT (primers 2 and 3), and those PCR fragments were replaced in the ΔEG2-5 construct through EcoRV/NotI sites. The introduction of EcoRV site in primers 1 and 2 did not produce any amino acid changes. ΔEG8-CT cDNA was amplified by PCR using primers 4 and 5. Constructs ΔEG6-7 and ΔEG8-9 were obtained by overlap extension. ^21^ Deletion of EG6-7 was introduced by primers 7 and 8, which were combined with primers 6 and 9, respectively, to generate the 5′ and 3′ fragments. The gel-purified fragments were fused by annealing overlapping ends and amplified with primers 6 and 9 to give the final product. The final product was inserted into full-length fibulin-1C (-stop codon) in pBluescript through BamHI/NheI sites. For the ΔEG8-9 construct, the 5′ fragment was generated by primers 6 and 10, and the 3′ fragment by primers 11 and 5. Then, those fragments were fused and amplified with primers 6 and 5, and inserted into the plasmid of fibulin-1C (-stop) in pBluescript through BamHI/NotI sites. All constructs were verified by Dye Terminator Cycle Sequencing (ABI/ThermoFisher Scientific, Waltham, MA, USA). Sodium dodecyl sulfate-polyacrylamide gel electrophoresis (SDS-PAGE) was performed for His-tagged fibulin-1C and recombinant proteins (Supplemental Figure 2).

Expi293 cells were transfected according to the manufacturer’s protocol (ThermoFisher Scientific); only deletion mutant ΔEG8-9 was not expressed. Harvested conditioned media were incubated with Talon Metal Affinity resin (Clontech, Silicon Valley, CA, USA) at 4 °C overnight, and His-tagged protein was eluted with 150 mM imidazole. Eluted proteins were dialyzed against 0.3 M NaCl/0.05 M Tris-HCl, pH 7.4. Recombinant fibulin-1C and -1D without the tags were prepared as described previously.^22^

### 2.6 Statistical analysis

ALL data are presented as mean ± standard deviation (SD). The Mann–Whitney test was used for analysis between two groups (*Figure 2, Figure 3, Figure 4E, Figure 5E, Figure 6, Supplemental Figure 3, Supplemental Figure 4, Supplemental Table 1, and Supplemental Table 3*). For comparisons among three or more groups, analysis was performed using the Kruskal–Wallis test followed by the Fisher least significant difference post hoc test and Mann–Whitney test (*Figure 4A, 4B, 4G, 4H, 4J, Figure 5B, and 5F*). A *P*-value < 0.05 was considered significant. All data were analyzed using GraphPad Prism software (GraphPad Software, La Jolla, CA, USA).

## 3. Results

### 3.1 EP4 was upregulated in the area of highly proliferative neointima

It has been demonstrated in wild-type mice that IH gradually develops after wire-induced vascular injury and reaches a maximum at 3 to 4 weeks. ^23^ Proliferating cell nuclear antigen (PCNA) immunostaining demonstrated that cell proliferation was detectable 1 week after injury, most prominent at 2 weeks after injury, and then decreased at 4 weeks (*Figure 1A*). To investigate the spatiotemporal EP4 expression in the injured artery, we used EP4 reporter mice (*Ptger4*-IRES-nlsLacZ). Expression of EP4 was found to be most abundant in the area of IH 2 weeks after vascular injury and decreased subsequently (*Figure 1B*), whereas X-gal positive signals were not observed in the injured C57BL/6N mice after vascular injury (*Figure 1C*). EP4-expressing cells did not demonstrate an immunoreaction for von Willebrand factor (a marker of endothelial cells), although protein expression of von Willebrand factor was detected in endothelial cells of the uninjured part as the arrows indicated in *Figure 1D*. CD68 (a marker of monocytes/macrophages) was expressed at the adventitial side of the injured artery, whereas EP4-expressing cells did not exhibit immunoreaction for CD68 (*Figure 1E*). These data suggested that EP4 was temporally and highly expressed in proliferative VSMCs in the region of IH after injury.

**Figure 1.**
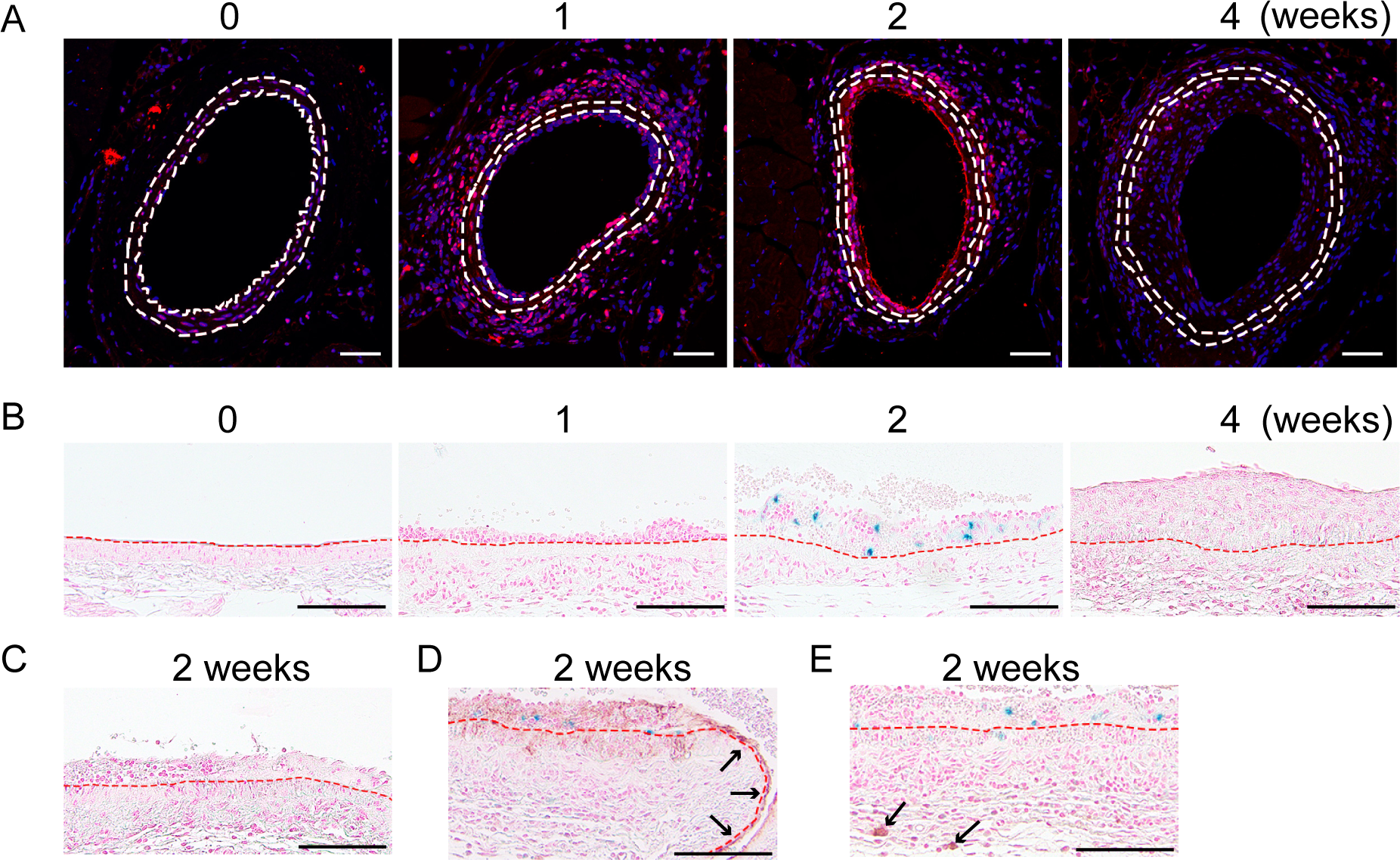
EP4 expression in the area of cell proliferative phase of IH. (*A*) Time course of representative images of immunofluorescence for PCNA in wire-injured femoral arteries in C57BL/6N mice. Scale bars represent 50 μm. (*B*) Time course of representative images of X-gal and Nuclear Fast Red staining of the long axis of wire-injured femoral arteries in *Ptger4*-IRES-nlsLacZ mice. Blue color indicates EP4 expression. (*C*) Representative images of X-gal and Nuclear Fast Red staining of the femoral artery 2 weeks after wire injury in C57BL/6N mice (negative control). (*D-E*) Representative images of immunohistochemistry for von Willebrand factor and CD68, which were stained with X-gal and Nuclear Fast Red, in C57BL/6N mouse femoral artery 2 weeks after vascular injury. Scale bars in *D-E* represent 100 μm. The internal and external elastic laminae are indicated by dotted lines. EP4, prostaglandin E_2_ receptor 4; IH, intimal hyperplasia; PCNA, proliferating cell nuclear antigen; X-gal, 4-chloro-5-bromo-3-indolyl-D-galactopyranoside; vWF, von Willebrand factor.

### 3.2 EP4 signaling in VSMCs exacerbated IH

We next investigated the role of EP4 using VSMC-selective deletion or overexpression EP4 mice. Because homozygous deficiency of EP4 in VSMCs causes neonatal death due to patent ductus arteriosus, ^24^ VSMC-selective heterozygous EP4 deficient mice, which have a 43% lower expression level of EP4 mRNA than *Ptger4*^+/+^;*SM22-Cre* mice^14^, were subjected to vascular injury.

The area of IH and the ratio of IH to the medial layer were significantly reduced in *Ptger4*^fl/+^;*SM22-Cre* mice compared to control mice (*Ptger4*^+/+^;*SM22-Cre*) at 2 and 4 weeks after vascular injury (*Figure 2A-F*). The area of the medial layer, area of internal lumen, length of the external elastic laminae, and thickness of the medial layer did not differ between *Ptger4*^fl/+^;*SM22-Cre* and controls (Supplemental Figure 3). There were no sex differences in injury-induced IH at 2 and 4 weeks after vascular injury (Supplemental Table 1).

**Figure 2.**
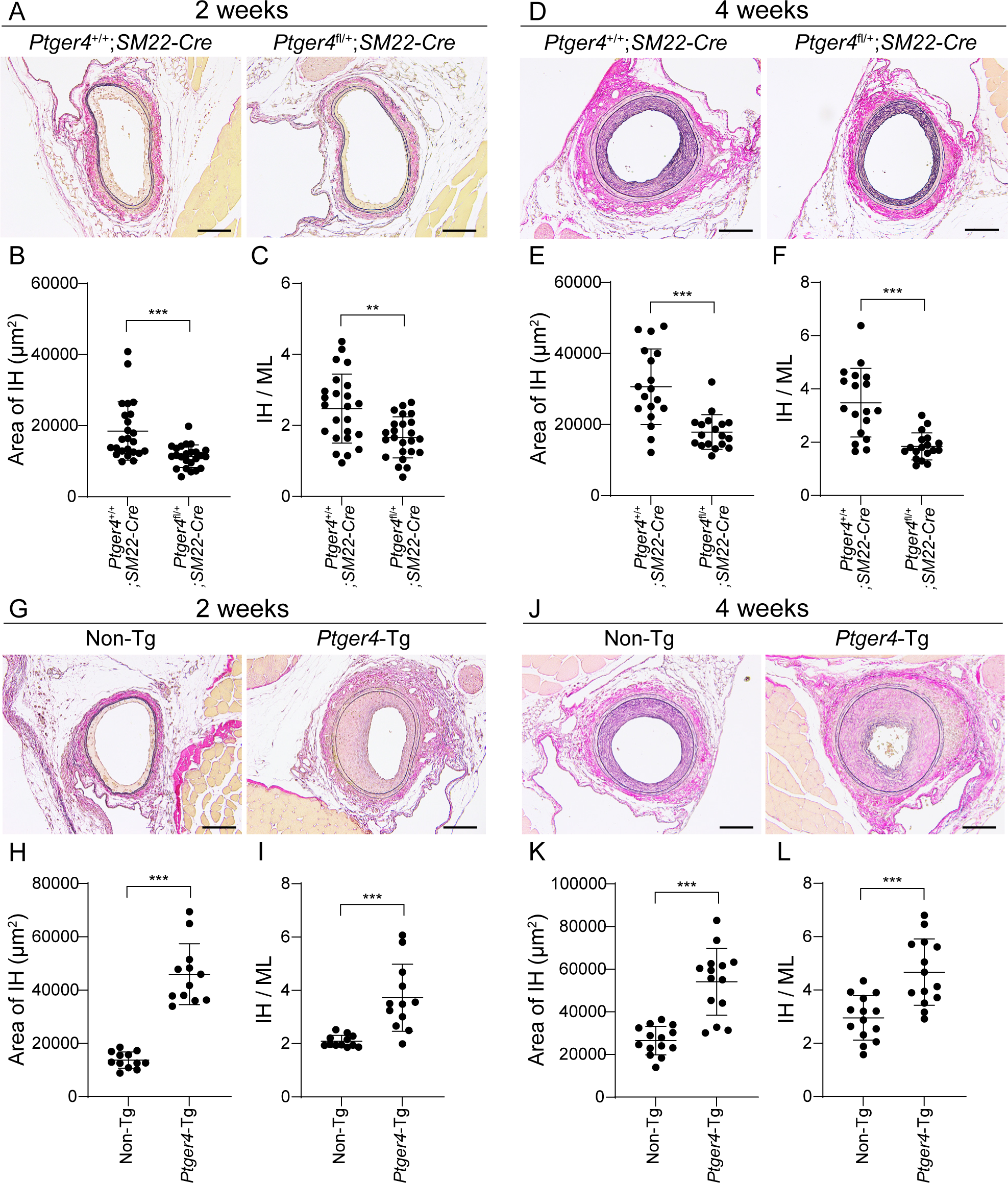
EP4 signaling increased injury-induced vascular IH. (*A*, *D*) Representative images of Elastica van Gieson staining of cross-sections of the femoral arteries at 2 and 4 weeks after injury in *Ptger4*^fl/+^;*SM22-Cre* and *Ptger4*^+/+^;*SM22-Cre* (control) mice. (*B*, *C*) Quantitative analysis of *A*; n = 24 (male: n = 13, female: n = 11). (*E*, *F*) Quantitative analysis of *D*; n = 18 (male: n = 9, female: n = 9). (*G*, *J*) Representative images of Elastica van Gieson staining of cross-sections of the femoral arteries 2 and 4 weeks after wire injury in non-Tg (control) and *Ptger4*-Tg mice. (*H*, *I*) Quantitative analysis of *G*; n = 12 (male: n = 6, female: n = 6). (*K*, *L*) Quantitative analysis of *J*; n = 14 (male: n = 7, female: n = 7). Scale bars represent 100 μm. \*\**P* < 0.01, \*\*\**P* < 0.001. IH, intimal hyperplasia; IH/ML, intimal hyperplastic area:medial layer area ratio; ML, medial layer.

Conversely, *Ptger4*-Tg mice exhibited markedly enhanced IH at 2 and 4 weeks following vascular injury compared to non-Tg control mice (*Figure 2G-L*). The area of medial layer, thickness of the medial layer, and length of the external elastic laminae were greater in *Ptger4*-Tg mice at 2 weeks after injury, and in *Ptger4*-Tg mice these parameters were equivalent to those of non-Tg mice 4 weeks later (Supplemental Figure 4). The area of internal lumen was significantly smaller in *Ptger4*-Tg mice than in non-Tg mice at 4 weeks post-injury (Supplemental Figure 4F). These data suggest that injury-induced EP4 stimulation in VSMCs confers a highly proliferative character.

### 3.3 Oral administration of an EP4 antagonist reduced IH

Based on the data showing that EP4 played a role in vascular IH, we investigated whether the blockade of EP4 signaling *in vivo* would reduce injury-induced IH. The dose of an EP4 antagonist (CJ42794) for *in vivo* experiments was determined based on a previous study. ^25^ Oral administration of CJ42794 during vascular injury attenuated the area of IH and the ratio of IH to the medial layer at 4 weeks after injury (*Figure 3*). These results using genetically, and pharmacologic approaches suggest that PGE_2_-EP4 signaling in VSMCs has a profound effect on promoting IH.

**Figure 3.**
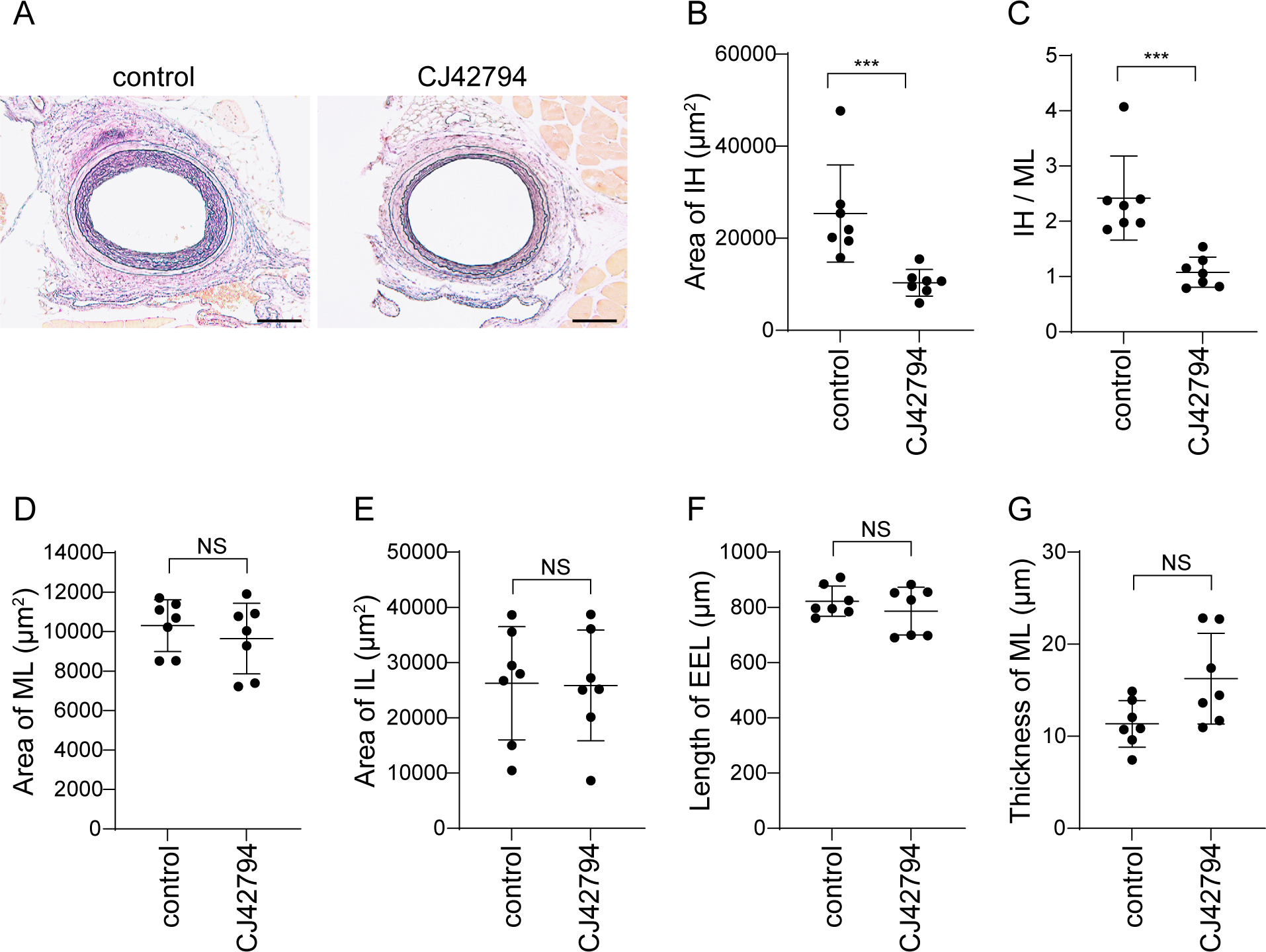
Oral administration of an EP4 antagonist reduced IH. (*A*) Representative images of Elastica van Gieson staining of cross-sections of femoral arteries 4 weeks after wire injury in C57BL/6N male mice treated orally twice daily with 0.5% methylcellulose (control) and CJ42794 (EP4 antagonist). Scale bars represent 100 μm. (*B*-*G*) Quantitative analysis of *A*; n = 7. \*\*\**P* < 0.001, NS: not significant. IH, intimal hyperplasia; IH/ML, intimal hyperplastic area:medial layer area ratio; ML, medial layer; IL, internal lumen; EEL, external elastic laminae.

### 3.4 PGE_2_-EP4 signaling increased fibulin-1 expression

To investigate the mechanisms by which EP4 exacerbated IH, we focused on the ECM. In a previous study, an unbiased expression analysis (accession number: GSE146758) demonstrated that PGE_2_ stimulation increased fibulin-1 in VSMCs isolated from *Ptger4*-Tg mice (*Ptger4*-Tg VSMCs). ^14^ Fibulin-1 is a secreted glycoprotein and associates with cardiovascular development^13, 26^ and proliferation in multiple cell types, including cancer cells and chondrocytes. ^27, 28^, Fibulin-1 has four splicing variants, A through D in humans. Mice, and presumably most other mammalian species, express only variants C and D, which are produced in comparable amounts in most tissues and cell lines studied. ^29^

Stimulation of *Ptger4*-Tg VSMCs with PGE_2_ increased both *Fbln1C* and *Fbln1D* mRNAs, which were decreased by PGE_2_ combined with an EP4 antagonist (*Figure 4A-B*). Western blotting demonstrated that EP4 stimulation increased secretion of fibulin-1 proteins in a dose- and time-dependent manner (*Figure 4C-E*).

**Figure 4.**
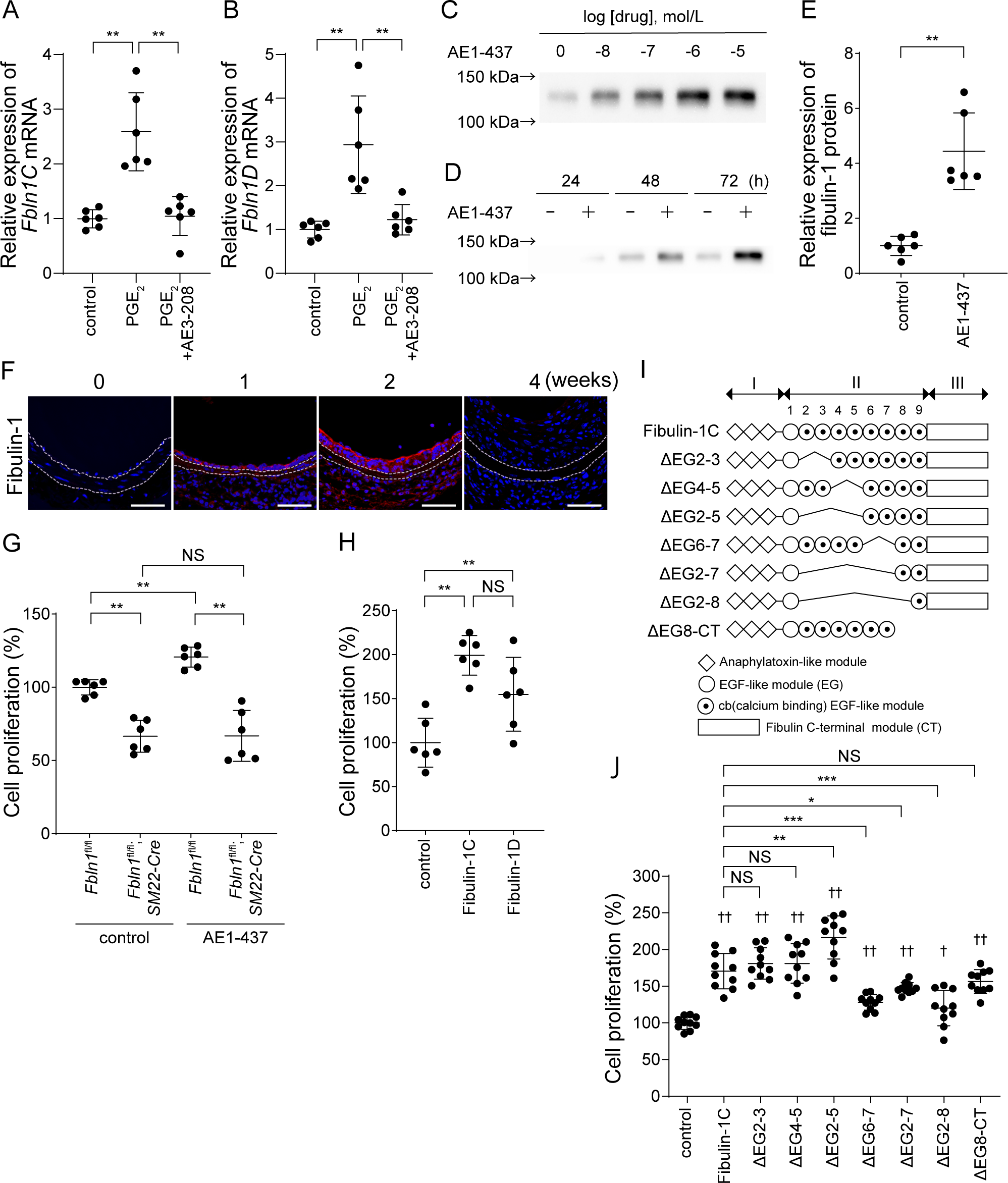
EP4-induced fibulin-1 promoted VSMC proliferation. (*A*, *B*) Expression of *Fbln1C* and *Fbln1D* mRNA in *Ptger4*-Tg VSMCs stimulated with PGE_2_ or PGE_2_ + ONO-AE3-208 (AE3-208, an EP4 antagonist); n = 6. (*C*, *D*) Concentration- and time course-dependent expression of fibulin-1 proteins in *Ptger4*-Tg VSMCs stimulated with ONO-AE1-437 (AE1-437, an EP4 agonist). (*E*) Quantitative analysis of expression of fibulin-1 protein by western blotting in *Ptger4*-Tg VSMCs stimulated with ONO-AE1-437 (AE1-437, 1 μmol/L) for 72 h; n = 6. (*F*) Time course of immunofluorescent staining for fibulin-1 (red) with Hoechst (blue) of cross-sections of wire-injured femoral arteries in C57BL/6N mice. The internal and external elastic laminae are indicated by dotted lines. Scale bars represent 50 μm. (*G*) Quantitative analysis of proliferation of *Fbln1*^fl/fl^ (control) and *Fbln1*^fl/fl^;*SM22-Cre* VSMCs after stimulation with ONO-AE1-437 (AE1-437, 1 μmol/L) for 48 h; n = 6. (*H*) The effect of fibulin-1C or -1D recombinant proteins (1 μg/mL, 48 h) on proliferation in non-Tg VSMCs; n = 6. (*I*) Structures of full-length and deletion mutants of fibulin-1C recombinant proteins. (*J*) Quantitative analysis of proliferation of non-Tg VSMCs after adding fibulin-1C recombinant proteins; n = 10. \**P* < 0.05, \*\**P* < 0.01, \*\*\**P* < 0.001, NS: not significant; †*P* < 0.05, ††*P* < 0.01 vs. controls.

We investigated the spatiotemporal expression of fibulin-1 proteins during vascular injury in wild-type mice and found that the immunofluorescent signal for fibulin-1 became visible in the area of IH at 1 week after vascular injury, and overt expression of fibulin-1 proteins in IH was observed at 2 weeks after injury (Figure 4F). Fibulin-1 expression was decreased 4 weeks after injury, similar to the expression pattern observed for EP4.

### 3.5 Fibulin-1 promoted VSMC proliferation

To examine whether fibulin-1 promotes cell proliferation, we used a primary culture of VSMCs isolated from *Fbln1*^fl/fl^;*SM22-Cre* mice and found that *Fbln1*-deficient VSMCs exhibited reduced proliferative ability compared to controls (*Figure 4G*). An EP4 agonist (ONO-AE1-437) increased proliferation in *Fbln1*^fl/fl^ VSMCs, whereas EP4 stimulation did not increase proliferation in *Fbln1*-deficient VSMCs (*Figure 4G*).

Recombinant proteins of fibulin-1C or -1D significantly promoted proliferation of non-Tg control VSMCs (*Figure 4H*). Although there was no difference in cell proliferation between the treatments of fibulin-1C and -1D, fibulin-1C recombinant protein showed greater stability and enhanced cell proliferation (*Figure 4H*). Therefore, we used fibulin-1C recombinant proteins in further experiments.

### 3.6 Calcium binding epithelial growth factor (cbEGF)-like module 6-8 of fibulin-1 contributed to cell proliferation

The fibulin family comprises of eight extracellular glycoproteins that characteristically share a globular domain at the carboxy terminus, which is called fibulin-type module, FC domain or domain III. This domain is preceded by a variable number of calcium binding epidermal growth factor (cbEGF)-like modules commonly referred to as domain II. The amino-terminal region of fibulins contains domain I. ^30^

Domain II of fibulin-1 comprises nine epidermal growth factor (EGF)-like modules, eight of which have consensus sequences for calcium binding. ^22^ The eight fibulin-1 calcium binding sites were suggested to maintain the structure and binding functions of fibulin-1. ^20^ To further investigate the mechanisms of fibulin-1–mediated cell proliferation, we generated multiple mutant recombinant proteins of fibulin-1 (*Figure 4I*).

Full-length fibulin-1C proteins increased proliferation in VSMCs (*Figure 4J*). The deletion of cbEGF-like module 2-3, 4-5, or 2-5 also increased VSMC proliferation, which were comparable in degree to that of full-length fibulin-1C. In contrast, the deletion of cbEGF-like module 6-7, 2-7, or 2-8 induced a significantly lesser proliferative response than full-length fibulin-1. The deletion of cbEGF-like module 8-9 and domain III seemed to result in loss of the proliferative effect, although this did not reach statistical significance. These data indicated that cbEGF-like module 6-7 or 6-8 contribute to VSMC proliferation.

### 3.7 ECM1 was upregulated by EP4 and expressed in IH

Fibulin-1 binds to multiple molecules and facilitates the organization of a variety of ECMs. ^31^ To investigate ECM proteins that potentially cooperate with fibulin-1, we examined the EP4-mediated changes in expression of 20 genes that reportedly bind to fibulin-1 using previous reported data (accession number: GSE146758) ^14^, as shown in Supplemental Table 2. The most upregulated gene was *Vcan*, but versican proteins were not detected at 2 weeks after vascular injury and were highly expressed in the area of IH at 4 weeks after injury (Supplemental Figure 5). We then focused on extracellular matrix protein 1 (ECM1) as the second most upregulated ECM protein and found that ECM1 was abundantly expressed in the area of IH at 2 weeks after vascular injury, which has similar expression pattern to EP4 and fibulin-1 (*Figure 5A*).

**Figure 5.**
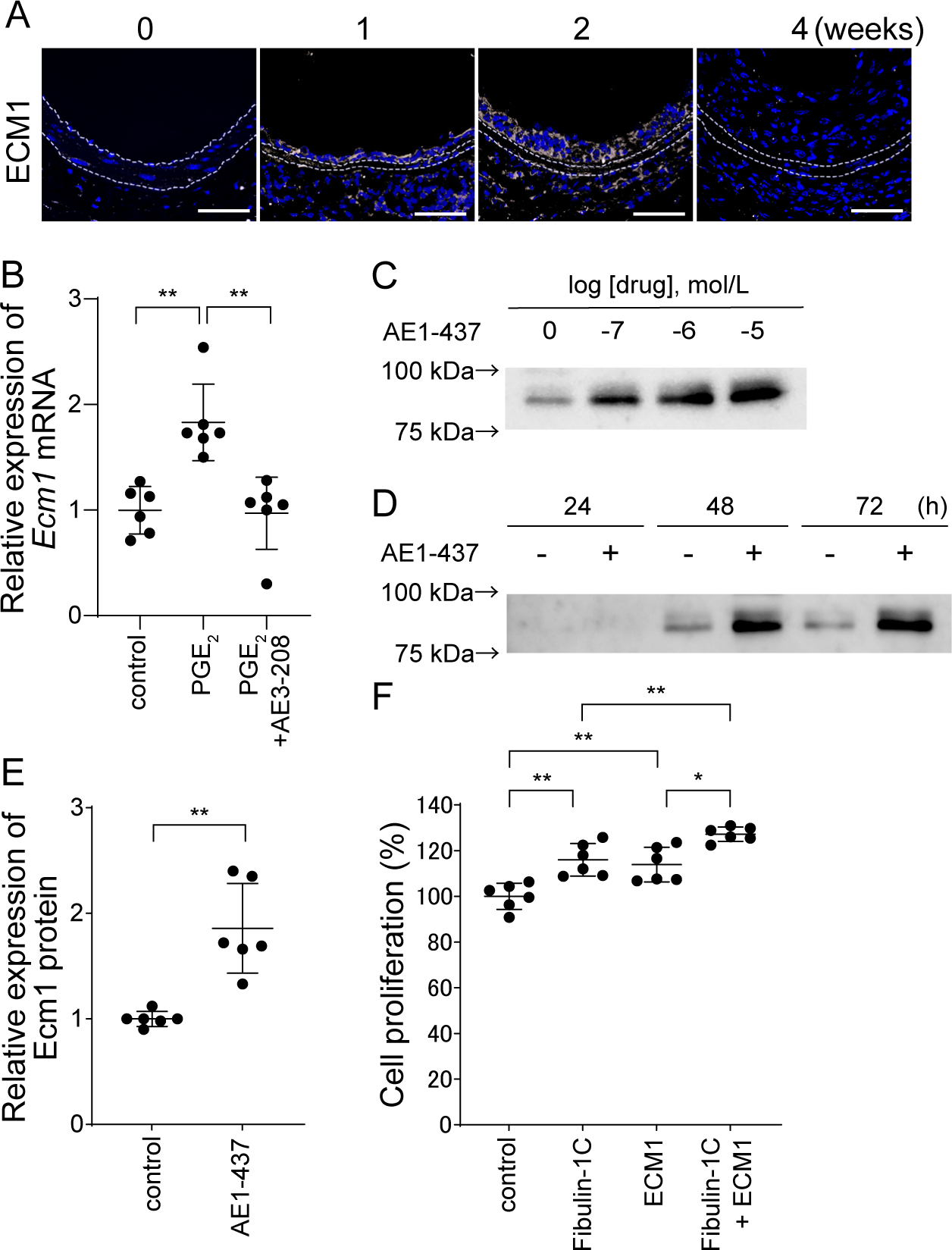
EP4-induced ECM1 promoted VSMC proliferation. (*A*) Time course of immunofluorescent staining for ECM1 (yellow) with Hoechst (blue) of cross-sections of wire-injured femoral arteries in C57BL/6N mice. The internal and external elastic laminae are indicated by dotted lines. Scale bars represent 50 μm. (*B*) Expression of *Ecm1* mRNA in *Ptger4*-Tg VSMCs stimulated with prostaglandin E_2_ (PGE_2_) and PGE_2_ + ONO-AE3-208 (AE3-208, an EP4 antagonist); n = 6. (*C*, *D*) Concentration- and time-course-dependent expression of ECM1 proteins in *Ptger4*-Tg VSMCs stimulated with ONO-AE1-437 (AE1-437, an EP4 agonist). (*E*) Quantitative analysis of expression levels of ECM1 protein determined by western blotting of *Ptger4*-Tg VSMCs stimulated with AE1-437 (1 μmol/L) for 72 h; n = 6. (*F*) Quantitative analysis of proliferation of non-Tg VSMCs after addition of fibulin-1C, ECM1, or fibulin-1C+ECM1; n = 6. \**P* < 0.05, \*\**P* < 0.01, NS: not significant.

### 3.8 ECM1 combined with fibulin-1 increased VSMC proliferation

ECM1 is an 85 kDa glycoprotein expressed throughout the body, ^32^ and it promotes proliferation of multiple types of cells, including endothelial cells and epidermal keratinocytes, and pancreatic ductal adenocarcinoma. ^33, 34^

The expression of *Ecm1* mRNA in *Ptger4*-Tg VSMCs increased following stimulation with PGE_2_, which was decreased by an EP4 antagonist (*Figure 5B*). Western blotting showed a concentration- and time-dependent increase in ECM1 after stimulation with an EP4 agonist (*Figure 5C-E*). Full-length fibulin-1C or ECM1 recombinant proteins similarly increased proliferation in non-Tg VSMCs, and ECM1 recombinant proteins enhanced fibulin-1C–mediated proliferation of VSMCs (*Figure 5F*).

### 3.9 Fibulin-1 deficiency attenuated IH

Finally, we investigated the contribution of fibulin-1 during vascular injury *in vivo*. Injury-induced IH (area of IH and ratio of IH to the medial layer) was reduced in the *Fbln1*^fl/fl^;*SM22-Cre* mice at 2 weeks post-injury compared to the control group (*Fbln1*^fl/fl^) (*Figure 6A-G*). Deficiency of fibulin-1 in VSMCs also attenuated IH 4 weeks later (*Figure 6H-N*). There were no sex differences in attenuation of injury-induced IH at 2 or 4 weeks after vascular injury (Supplemental Table 3).

**Figure 6.**
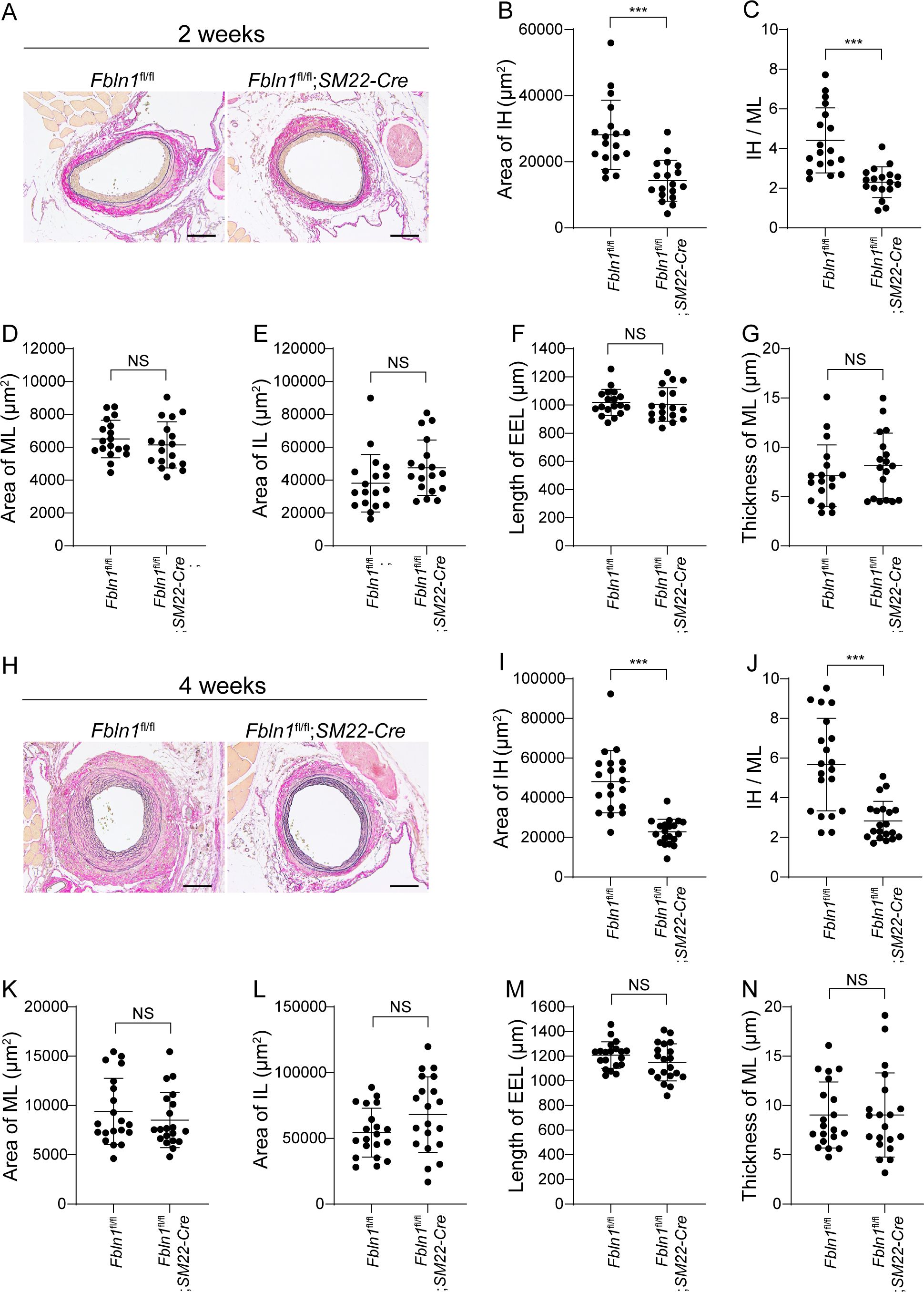
Deficiency of fibulin-1 attenuated injury-induced vascular IH. (*A*, *H*) Representative images of Elastica van Gieson staining of cross-sections of the femoral arteries 2 and 4 weeks after wire injury of *Fbln1*^fl/fl^;*SM22-Cre* and *Fbln1*^fl/fl^ (control) mice. Scale bars represent 100 μm. (*B*-*G*) Quantitative analysis of *A*; n = 18 (male: n = 9, female: n = 9). (*I*-*N*) Quantitative analysis of *H*; n = 20 (male: n = 10, female: n = 10). \*\*\**P* < 0.001, NS: not significant. IH, intimal hyperplasia; IH/ML, intimal hyperplastic area:medial layer area ratio; ML, medial layer; IL, internal lumen; EEL, external elastic laminae.

## 4. Discussion

Our results demonstrated that the PGE_2_–EP4–fibulin-1 axis in VSMCs increased IH through cell proliferation. The fibulin-1 cbEGF-like modules 6-7 or 6-8 were associated with VSMC proliferation, and fibulin-1 and its binding partner ECM1 additively increased VSMC proliferation and exacerbated IH.

EP4 is expressed in a variety of cell types; it maintains homeostasis in physiological conditions and exacerbates or ameliorates in pathological conditions. ^35^ In injured vessels, VSMCs, endothelial cells, and immune infiltrates play roles in IH, and these cell types express EP4 to some degree. It has been reported that EP4 signaling in endothelial cells and VSMCs is protective and destructive, respectively, against vascular injuty. ^9, 10^ Although cell type–specific roles of EP4 in IH have been indicated, the spatiotemporal EP4 expression pattern has not been fully understood because the specificity of anti-EP4 antibody can be compromised in mouse specimens. Activation of EP4 is implicated to promote EP4 expression in a positive feedback loop. ^36, 37^ Therefore, we generated *Ptger4*-IRES-nlsLacZ mice in which EP4 expression could be monitored without interference from EP4 signaling. To the best of our knowledge, this is the first study to show the spatiotemporal expression of EP4 during vascular injury. We demonstrate that EP4 is highly expressed in VSMCs in the area of IH. X-gal staining signals for EP4 in endothelial cells and CD68-positive cells did not reach the level of visualization. The use of smooth muscle–specific EP4 knockdown and EP4 overexpression mice and pharmacological inhibition of EP4 signaling indicated that EP4 in VSMCs exacerbated injury-induced IH. These data emphasize the importance of EP4 signaling in VSMCs in IH.

ECMs, which account for more than half the volume of human IH regions, ^38^ are major factors in restenosis in humans, a process that can occur months to years after stenting or balloon angioplasty. ^39, 40^ It has been demonstrated that neointimal versican and hyaluronan accumulation is a relatively early event after vascular injury, and late neointimal ECM contraction is associated with the replacement of water-trapping proteoglycans and hyaluronan with type I collagen. ^39, 40^ In the present study, we identified fibulin-1 as a potential positive driver of IH. Versican was abundantly expressed in IH 4 weeks after vascular injury, and fibulin-1 expression preceded versican expression. Together with the findings of PCNA staining, these data suggest that fibulin-1 contributes to the early neointimal proliferative phase.

Fibulin-1 is a secreted glycoprotein that belongs to the fibulin family. ^31^ Recently, fibulin-1 has gained attention in the research field of smooth muscle and cardiovascular disease. The levels of circulating fibulin-1 are positively correlated with arterial stiffness. ^41, 42^ Fibulin-1 is indicated to regulate tissue remodeling via promotion of cell migration in the heart and ductus arteriosus^13, 26^ There are four splice variants of fibulin-1 in humans. Variants C and D are expressed to the same extent in most tissues and cell lines, whereas variants A and B are only detected at low levels in human placenta. ^29^ Fibulin-1 consists of N-terminal (domain I) and C-terminal (domain III) globular structures connected by a central rod (domain II) composed of nine epidermal growth factor modules, eight of which possess a consensus sequence for calcium binding. The differences between fibulin-1C and -1D lie within in domain III in which fibulin-1C has a shorter domain III. ^22^

Fibulin-1 mediates various types of intracellular signals by binding to ECM proteins such as fibronectin, versican, aggrecan, and nidogen. ^31^ Most of these ECM proteins were demonstrated to bind to cbEGF-like modules of fibulin-1. ^22, 43, 44^ Electron microscopy reveals that full-length fibulin-1 has a dumbbell structure with a length of 32 nm. When the cbEGF-like modules 2-8 are removed, the rod-like structure is absent, and deletion of cbEGF-like modules 2-5 shortens the rod-like structure to 10 nm, ^20^ indicating that cbEGF-like modules are critical for the structural integrity and function of fibulin-1. ^45^ Our study demonstrated that the proliferative ability of VSMC was reduced when cbEGF-like modules 6-8 were deficient but not when cbEGF-like modules 2-5 were deleted. Based on these data, we can speculate that the specific function of cbEGF-like modules 6-8 or the integrity of rod-like structure might be important for VSMC proliferation. Although several studies have demonstrated the involvement of fibulin-1 in cell proliferation, the molecular mechanisms by which cells transduce pericellular fibulin-1 proteins to intracellular signals are largely unknown. We used several inhibitors to investigate how fibulin-1 promotes cell proliferation, but mitogen-activated protein kinase (MAPK) pathways and signal transducer and activator of transcription 3 (STAT3) were not involved. Further study is required to explore the receptor for fibulin-1 and its intracellular signaling pathway.

Like fibulin-1, ECM1 is a secreted glycoprotein. ^32^ ECM1 contains a 19-amino acid residue signal peptide, an N-terminal cysteine-free domain, two tandem repeat domains, and a C-terminal domain. ^33^ The latter three domains contain the characteristic cysteine arrangement CC-(X7-10)-C pattern, which generates double-loop structures for protein-protein interactions. ^33^ ECM-1 interacts with many other ECM proteins, including fibulin-1, fibulin-3, a proteoglycan perlecan, cartilage oligomeric matrix protein, and type IV collagen. ^33^ Domain III of fibulin-1C and -1D binds to the second cysteine-rich tandem repeat domain. ^46^

The biological function of ECM1 in blood vessels has been mainly studied in endothelial cells, because the skin pathology of patients with the genodermatosis lipoid proteinosis and lichen sclerosus, which genetically and immunologically target ECM1, respectively, lacks a normal capillary network in the upper dermis and enlargement of the dermal blood vessels. ^33^ It was reported that recombinant human ECM1 stimulated the proliferation of cultured endothelial cells and promotes the formation of blood vessels in the chorioallantoic membrane of chicken embryos, suggesting a functional role in angiogenesis. ^47^ In addition to the role of ECM1 in endothelial cells, the present study revealed that ECM1 was secreted by EP4 signaling in VSMCs and promoted cell proliferation. Similar to our results showing simultaneous upregulation of fibulin-1 and ECM1, a recent study demonstrated that ECM1 expression is associated with fibulin-1 upregulation in mouse myocardial infarction–induced cardiac fibrosis. ^48^ That study showed that ECM1 bound to low density lipoprotein receptor-related protein 1 (LRP1) and stimulated the fibroblast-to-myofibroblast transition. ^48^ LRP is reportedly involved in VSMC proliferation. ^49^ However, the molecular mechanisms of ECM1-mediated cell proliferation still need to be elucidated. In addition, exploring the cooperative roles of fibulin-1 and ECM1 is an important area for future investigations.

The present study showed no sex difference in the proliferative role of EP4 and fibulin-1 in IH, and an EP4 antagonist targeting pathologically upregulated EP4 is a potential therapeutic strategy against injury-induced IH. An EP4 antagonist has been examined up to phase 2 in a US pharmacological study (ClinicalTrials.gov ID: NCT03163966) and found to be nontoxic. In contrast, an EP4 agonist attenuated wire injury-induced IH through stimulating endothelialization. ^8^ Together with previous reports, ^12, 50^ these findings suggest that EP4 signaling at baseline or a moderate level is required for vascular development and physiology. Therefore, defining a dosage of an EP4 antagonist that attenuates excessive EP4 signaling but does not disrupt homeostasis is critical. In addition to regulation of EP4 signaling, the ECM should be considered a potential target for new approaches to prevent IH. Although the regulation and metabolism of fibulin-1 are largely unknown, targeting fibulin-1 may be a therapeutic strategy for IH by inhibiting fibulin-1–mediated VSMC proliferation in the early phase of IH, i.e., a decrease in fibulin-1 expression, degradation of fibulin-1, or blocking fibulin-1–mediated cellular signaling.

### 4.1 Conclusions

PGE_2_-EP4 signaling was activated in injury-induced neointima, and EP4-induced fibulin-1 cooperated with ECM1 to promote IH through VSMC proliferation. The cbEGF-like modules 6-7 or 6-8 of fibulin-1 were indicated to contribute to cell proliferation.

## Data Availability Statement

The data that support the findings of this study are available upon request from the corresponding author.

## Acknowledgments

The authors are grateful to Yuka Sawada for technical assistance.

