## supplementals and materials for "Spatiotemporal EP4-fibulin-1 expression is associated with vascular intimal hyperplasia"

Short title: **Fibulin-1 promotes intimal hyperplasia**

Shigekuni Okumura[1,2]; Sayuki Oka[1]; Takako Sasaki[3]; Marion A. Cooley[4]; Yuko Hidaka[1]; Shota Tanifuji[1]; Mari Kaneko[5]; Takaya Abe[5]; Richard M. Breyer[6]; Hiroshi Homma[2]; Yuko Kato[1,7]; Utako Yokoyama[1]

Affiliations:
[1]Department of Physiology, Tokyo Medical University, Tokyo, Japan
[2] Department of Emergency and Critical Care Medicine, Tokyo Medical University, Tokyo, Japan
[3]Department of Pharmacology, Faculty of Medicine, Oita University, Oita, Japan
[4]Department of Oral Biology and Diagnostic Sciences, Augusta University, GA, USA
[5]Laboratory for Animal Resources and Genetic Engineering, RIKEN Center for Biosystems Dynamics Research, Kobe, Japan
[6]Department of Medicine, Vanderbilt University Medical Center, Nashville, TN, USA
[7]Department of Advanced Medical Science, Faculty of Medicine, Oita University, Oita, Japan

Supplemental Figure 1

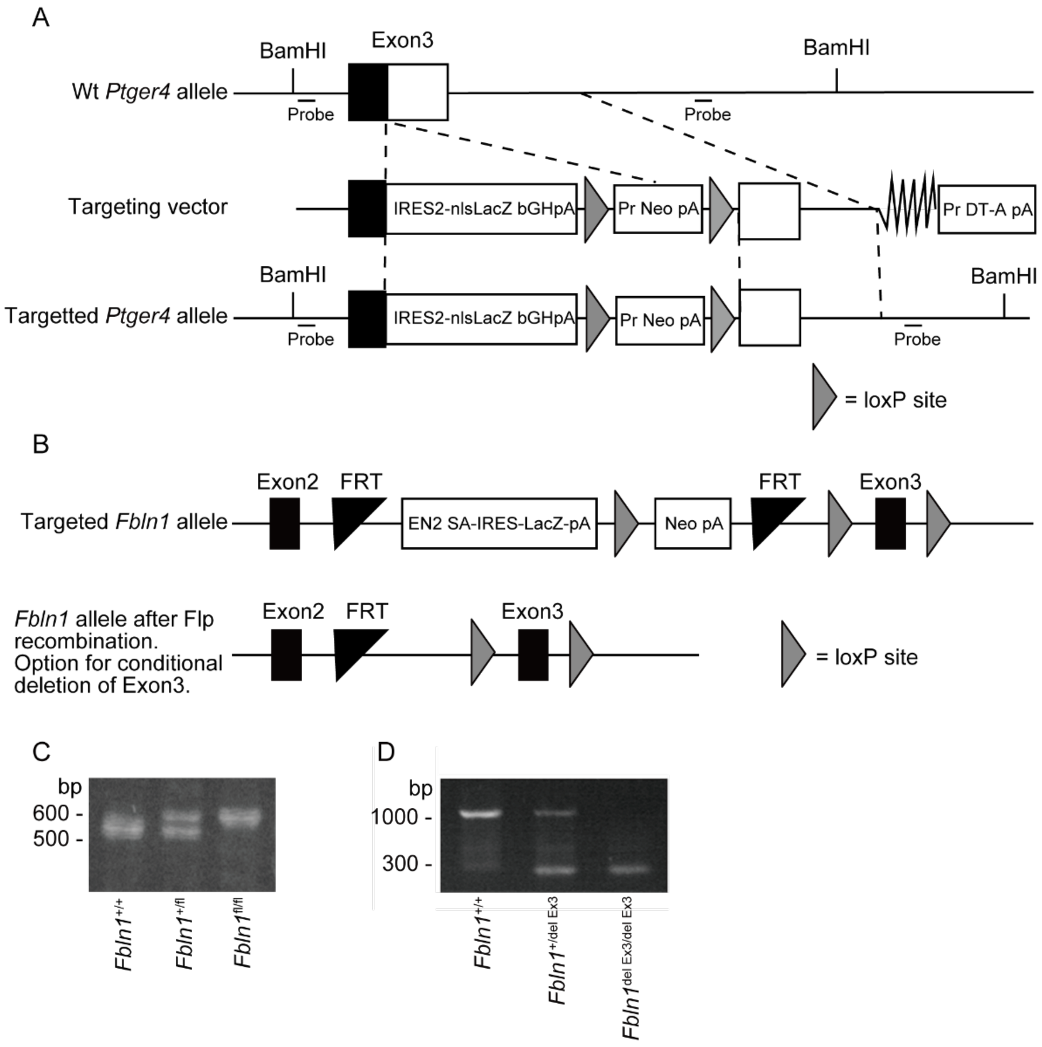


**Supplemental Figure 1.** Construction of *Ptger4*-IRES-nlsLacZ and $Fbln1^{fl/fl}$ mice. (*A*) *Ptger4*-IRES-nlsLacZ was produced by homologous recombination using ES cells. IRES: internal ribosomal entry site-nuclear localization signal, nls: nuclear localization signal, bGHpA: bovine growth hormone polyadenylation, Neo: neomycin resistance gene, Pr: promotor, pA: poly A. (*B*) *Fbln1* allele following homologous recombination with the targeting vector (upper panel). *Fbln1* allele after flipase excision of β-galactosidase (LacZ) and neomycin resistance (Neo) FRT flanked genes (lower panel). Following recombination, the resulting allele gives the option for conditional *Fbln1* deletion. (*C*) PCR genotyping of DNA from a wild-type, $Fbln1^{+/fl}$ and $Fbln1^{fl/fl}$ mouse. (*D*) PCR genotyping of DNA from a wild-type mouse and mice with deletion of exon 3 on one floxed allele ($Fbln1^{+/del\ Ex3}$) or both *Fbln1* floxed alleles ($Fbln1^{del\ Ex3/del\ Ex3}$).

Supplemental Figure 2

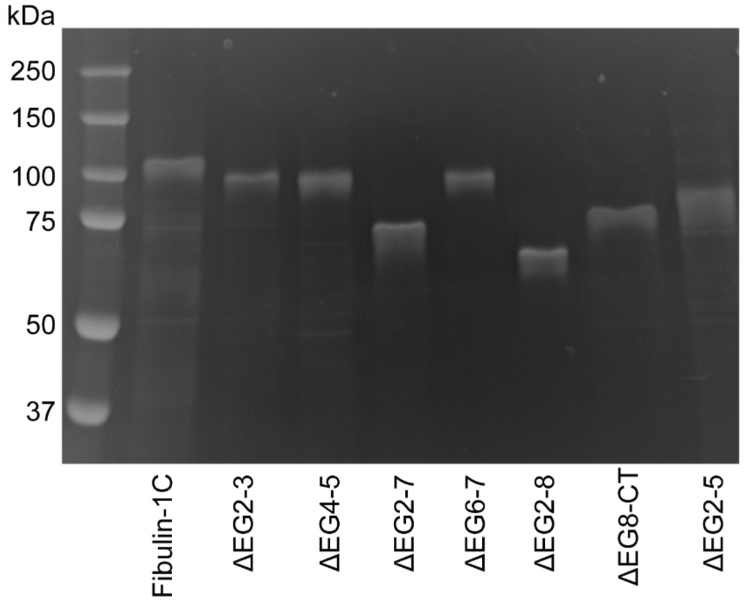


**Supplemental Figure 2.** SDS-PAGE performed on His-tagged fibulin-1C proteins. 1 µg of each protein was loaded and electrophoresis was performed under reducing conditions.

Supplemental Figure 3

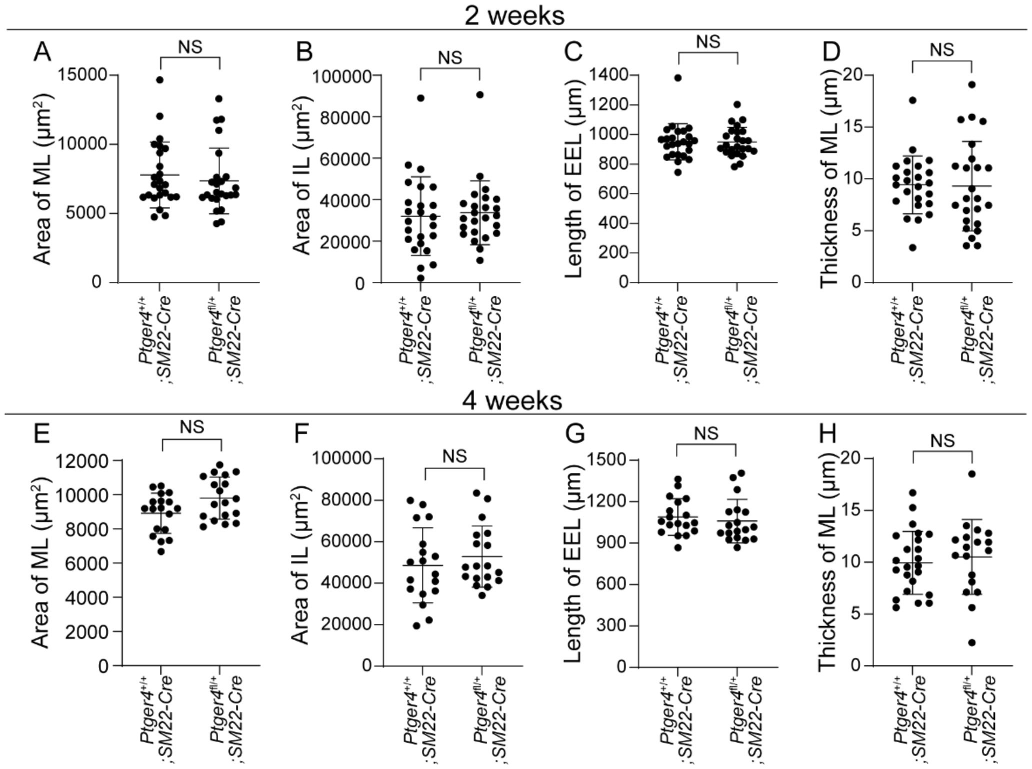


**Supplemental Figure 3.** EP4 increased intimal hyperplasia through VSMC. (*A-D*) Quantitative analysis of cross sections of femoral arteries 2 weeks after wire injury in *Ptger4*$^{fl/+}$;*SM22-Cre* and *Ptger4*$^{+/+}$;*SM22-Cre* (control) mice; n = 24 (male: n = 13, female: n=11). (*E-H*) Quantitative analysis of cross sections of femoral arteries 4 weeks after wire injury in *Ptger4*$^{fl/+}$;*SM22-Cre* and *Ptger4*$^{+/+}$;*SM22-Cre* (control) mice; n = 18 (male: n = 9, female: n = 9). ML, medial layer; IL, internal lumen; EEL, external elastic laminae. NS: not significant.

Supplemental Figure 4

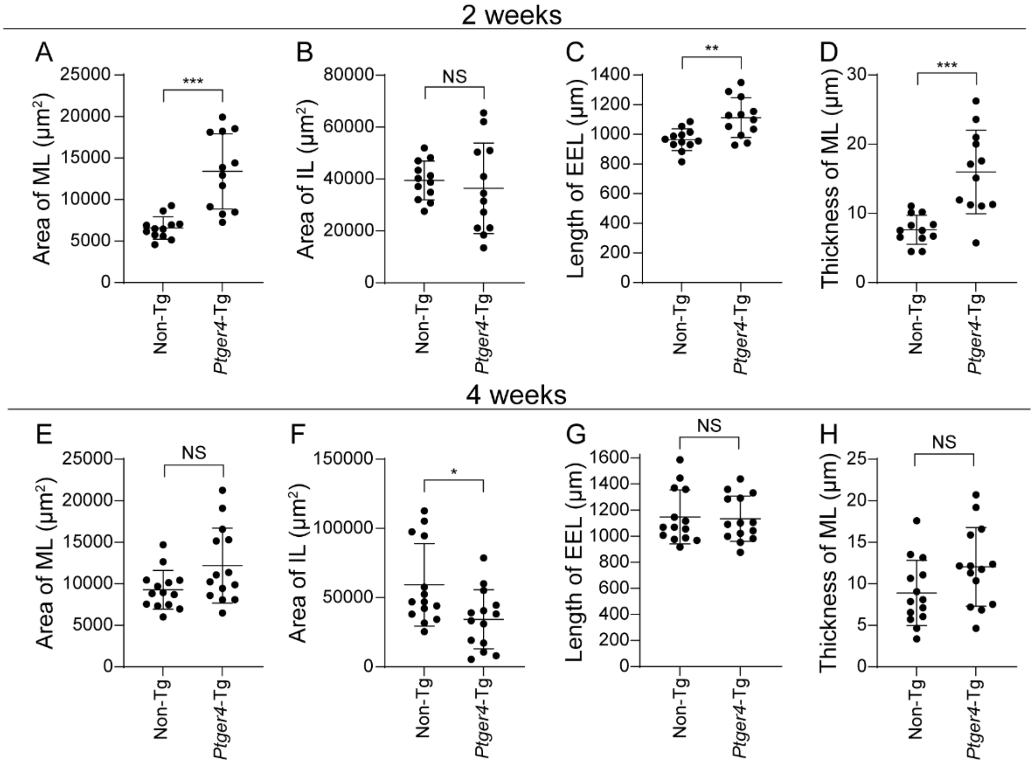


**Supplemental Figure 4.** EP4 increased intimal hyperplasia through VSMCs. (*A-D*) Quantitative analysis of cross sections of femoral arteries 2 weeks after wire injury in *Ptger4*-Tg and non-Tg (control) mice; n = 12 (male: n = 6, female: n = 6). (*E-H*) Quantitative analysis of cross sections of femoral arteries 4 weeks after wire injury in *Ptger4*-Tg and non-Tg (control) mice; n = 14 (male: n = 7, female: n = 7). ML, medial layer; IL, internal lumen; EEL, external elastic laminae. *$P < 0.05$, **$P < 0.01$, ***$P < 0.001$, NS: not significant.

Supplemental Figure 5

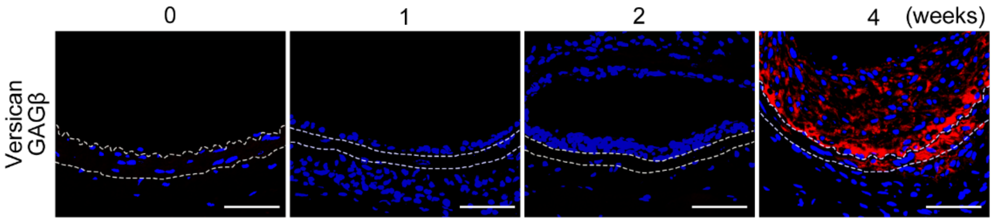


**Supplemental Figure 5.** Versican GAGβ different from EP4 and fibulin-1, increased 4 weeks after vascular injury. Time course of immunofluorescent staining for versican GAGβ (red) with Hoechst (blue) of cross sections of wire-injured femoral arteries in C57BL/6N mice. The internal and external elastic laminae are indicated by dotted lines. Scale bars represent 50 μm.

**Supplemental Table 1**. Intimal hyperplasia (IH) after vascular injury in vascular smooth muscle-selective heterozygous deficiency of *Ptger4* mice

| | Male (n = 13) 2 weeks after injury | | | | | | |
|---|---|---|---|---|---|---|---|
| | *Ptger4*$^{+/+}$;*SM22-Cre* | | | *Ptger4*$^{fl/+}$;*SM22-Cre* | | | *P*-value |
| Area of IH (μm$^2$) | 18,191 | ± | 10,097 | 11,315 | ± | 3762 | 0.019* |
| IH/ML | 2.2 | ± | 0.8 | 1.6 | ± | 0.4 | 0.044* |
| Area of IL (μm$^2$) | 39,726 | ± | 20,659 | 36,846 | ± | 20,116 | 0.579 |
| Area of ML (μm$^2$) | 8456 | ± | 2751 | 7317 | ± | 1924 | 0.336 |
| Length of EEL (μm) | 1003 | ± | 129 | 1002 | ± | 93 | 0.920 |
| Thickness of ML (μm) | 10.1 | ± | 3.1 | 10.4 | ± | 4.6 | 0.870 |
| | Female (n = 11) 2 weeks after injury | | | | | | |
| | *Ptger4*$^{+/+}$;*SM22-Cre* | | | *Ptger4*$^{fl/+}$;*SM22-Cre* | | | *P*-value |
| Area of IH (μm$^2$) | 18,854 | ± | 5733 | 11,462 | ± | 2497 | 0.002** |
| IH/ML | 2.9 | ± | 1.1 | 1.8 | ± | 0.7 | 0.007** |
| Area of IL (μm$^2$) | 22,986 | ± | 12,377 | 30,043 | ± | 5834 | 0.088 |
| Area of ML (μm$^2$) | 7011 | ± | 1667 | 7426 | ± | 2933 | >0.999 |
| Length of EEL (μm) | 889 | ± | 79 | 887.4 | ± | 69 | >0.999 |
| Thickness of ML (μm) | 8.7 | ± | 2.2 | 8.0 | ± | 3.7 | 0.216 |
| | Male (n = 9) 4 weeks after injury | | | | | | |
| | *Ptger4*$^{+/+}$;*SM22-Cre* | | | *Ptger4*$^{fl/+}$;*SM22-Cre* | | | *P*-value |
| Area of IH (μm$^2$) | 38,236 | ± | 8404 | 19,507 | ± | 5965 | 0.001*** |
| IH/ML | 4.5 | ± | 0.9 | 2.0 | ± | 0.7 | <0.0001*** |
| Area of IL (μm$^2$) | 48,610 | ± | 18,746 | 52,200 | ± | 18,866 | 0.730 |
| Area of ML (μm$^2$) | 8489 | ± | 1158 | 9958 | ± | 1186 | 0.040* |
| Length of EEL (μm) | 1130 | ± | 122 | 1062 | ± | 201 | 0.190 |
| Thickness of ML (μm) | 9.0 | ± | 2.1 | 10.3 | ± | 2.5 | 0.287 |
| | Female (n = 9) 4 weeks after injury | | | | | | |
| | *Ptger4*$^{+/+}$;*SM22-Cre* | | | *Ptger4*$^{fl/+}$;*SM22-Cre* | | | *P*-value |
| Area of IH (μm$^2$) | 22,992 | ± | 6273 | 16,290 | ± | 3087 | 0.024* |
| IH/ML | 2.5 | ± | 0.7 | 1.7 | ± | 0.2 | 0.011* |
| Area of IL (μm$^2$) | 48,711 | ± | 18,680 | 53,563 | ± | 10,285 | 0.489 |
| Area of ML (μm$^2$) | 9334 | ± | 1083 | 9649 | ± | 1321 | 0.667 |
| Length of EEL (μm) | 1049 | ± | 138 | 1057 | ± | 109 | 0.796 |
| Thickness of ML (μm) | 10.5 | ± | 3.4 | 10.7 | ± | 4.6 | 0.988 |

IH, intimal hyperplasia; IH/ML, intimal hyperplasic area:medial layer area ratio; ML, medial layer; IL, internal lumen; EEL, external elastic laminae.**P* < 0.05, ***P* < 0.01,

***$P < 0.001$.

**Supplemental Table 2**. EP4 signaling–mediated mRNA expression changes in fibulin-1–binding partners.

| GenBank Accession | Description | Average of control group | Average of $PGE_2$ stimulation group | Fold change $PGE_2$/ control |
|---|---|---|---|---|
| NM_001081249.1 | Versican (*Vcan*) | 363.9 | 16,601.3 | 45.6 |
| NM_181849.3 | Fibrinogen beta chain (*Fgb*) | 8.9 | 62.3 | 7.0 |
| NM_007899.5 | Extracellular matrix protein 1 (*Ecm1*) | 41,074.2 | 127,015.5 | 3.1 |
| NM_177544.4 | Angiogenin4 (*Ang4*) | 50.9 | 144.0 | 2.8 |
| NM_001011876.2 | Angiogenin6 (*Ang6*) | 135.3 | 307.9 | 2.3 |
| NM_001123394.2 | Angiogenin3 (*Ang3*) | 64.2 | 116.0 | 1.8 |
| NM_007449.3 | Angiogenin2 (*Ang2*) | 31.7 | 49.3 | 1.6 |
| NM_001161731.3 | Angiogenin (*Ang*) | 187.0 | 288.7 | 1.5 |
| NM_009621.5 | ADAM metallopeptidase with thrombospondin type 1 motif 1 (*Adamts1*) | 13,714.1 | 13,182.6 | 1.0 |
| NM_001198823.1 | Amyloid beta precursor protein (*App*) | 15,020.9 | 14,833.3 | 1.0 |
| NM_001276408.1 | Fibronectin 1(*Fn1*) | 146,943.6 | 136,167.7 | 0.9 |
| NM_010415.2 | Heparin binding EGF like growth factor (*Hbegf*) | 1050.9 | 995.7 | 0.9 |
| NM_011367.2 | Sex hormone binding globulin (*Shbg*) | 8.9 | 7.5 | 0.8 |
| NM_010917.3 | Nidogen1 (*Nid1*) | 2192.8 | 1416.9 | 0.6 |
| NM_008480.2 | Laminin subunit alpha 1 (*Lama1*) | 54.4 | 34.9 | 0.6 |
| NM_010930.5 | Cellular communication network factor 3 (*Ccn3*) | 2881.2 | 1064.1 | 0.4 |
| NM_001361500.1 | Aggrecan (*Acan*) | 40.5 | 13.3 | 0.3 |
| NM_008481.2 | Laminin subunit alpha 2 (*Lama2*) | 307.1 | 94.5 | 0.3 |
| NM_024237.4 | Fibulin 7 (*Fbln7*) | 687.5 | 116.2 | 0.2 |
| NM_007925.4 | Elastin (*Eln*) | 858.9 | 9.0 | 0.0 |

EP4-Tg VSMCs were stimulated with $PGE_2$ for 24 h.

**Supplemental Table 3** Intimal hyperplasia (IH) after vascular injury in vascular smooth muscle-selective *Fibulin1*-deficient mice

| | Male (n = 9) 2 weeks after injury | | | | | | |
|---|---|---|---|---|---|---|---|
| | *Fbln1*$^{fl/fl}$ | | | *Fbln1*$^{fl/fl}$;*SM22-Cre* | | | *P*-value |
| Area of IH (μm$^2$) | 27,019 | ± | 8601 | 12,955 | ± | 4916 | 0.003** |
| IH/ML | 4.4 | ± | 1.6 | 2.1 | ± | 0.7 | 0.003** |
| Area of IL (μm$^2$) | 34,369 | ± | 11,506 | 42,580 | ± | 7984 | 0.136 |
| Area of ML (μm$^2$) | 6301 | ± | 1016 | 5997 | ± | 1071 | 0.605 |
| Length of EEL (μm) | 982 | ± | 62 | 952 | ± | 60 | 0.258 |
| Thickness of ML (μm) | 7.4 | ± | 4.0 | 8.6 | ± | 3.3 | 0.328 |
| | Female (n = 9) 2 weeks after injury | | | | | | |
| | *Fbln1*$^{fl/fl}$ | | | *Fbln1*$^{fl/fl}$;*SM22-Cre* | | | *P*-value |
| Area of IH (μm$^2$) | 29,292 | ± | 12,456 | 15,620 | ± | 7325 | 0.011* |
| IH/ML | 4.5 | ± | 1.8 | 2.5 | ± | 0.8 | 0.003** |
| Area of IL (μm$^2$) | 41,989 | ± | 22,004 | 52,554 | ± | 22,006 | 0.222 |
| Area of ML (μm$^2$) | 6702 | ± | 1280 | 6299 | ± | 1732 | 0.546 |
| Length of EEL (μm) | 1057 | ± | 105 | 1056 | ± | 143 | 0.931 |
| Thickness of ML (μm) | 6.8 | ± | 2.3 | 7.7 | ± | 3.5 | 0.716 |
| | Male (n = 10) 4 weeks after injury | | | | | | |
| | *Fbln1*$^{fl/fl}$ | | | *Fbln1*$^{fl/fl}$;*SM22-Cre* | | | *P*-value |
| Area of IH (μm$^2$) | 54,949 | ± | 16,969 | 23,159 | ± | 5118 | <0.0001*** |
| IH/ML | 6.1 | ± | 2.7 | 2.7 | ± | 1.0 | 0.005** |
| Area of IL (μm$^2$) | 44,556 | ± | 9074 | 65,400 | ± | 9074 | 0.089 |
| Area of ML (μm$^2$) | 10,224 | ± | 3433 | 9130 | ± | 2504 | 0.481 |
| Length of EEL (μm) | 1191 | ± | 90 | 1140 | ± | 144 | 0.280 |
| Thickness of ML (μm) | 9.8 | ± | 3.9 | 9.7 | ± | 4.2 | 0.897 |
| | Female (n = 10) 4 weeks after injury | | | | | | |
| | *Fbln1*$^{fl/fl}$ | | | *Fbln1*$^{fl/fl}$;*SM22-Cre* | | | *P*-value |
| Area of IH (μm$^2$) | 41,263 | ± | 11,374 | 22,395 | ± | 7671 | 0.001*** |
| IH/ML | 5.2 | ± | 1.9 | 3.0 | ± | 1.0 | 0.007** |
| Area of IL (μm$^2$) | 64,227 | ± | 20,766 | 70,875 | ± | 30,944 | 0.436 |
| Area of ML (μm$^2$) | 8548 | ± | 3258 | 7912 | ± | 3070 | 0.739 |
| Length of EEL (μm) | 1228 | ± | 124 | 1160 | ± | 164 | 0.393 |
| Thickness of ML (μm) | 8.3 | ± | 2.6 | 8.4 | ± | 4.5 | 0.698 |

IH, intimal hyperplasia; IH/ML, intimal hyperplasic area:medial layer area ratio; ML, medial layer; IL, internal lumen; EEL, external elastic laminae.*$P < 0.05$, **$P < 0.01$,

***$P < 0.001$.

**Supplemental Table 4**. Primary antibodies for immunofluorescent staining and western blotting

| Target antigen | Vendor or Source | Catalog # | Working concentration | Lot # |
|---|---|---|---|---|
| Fibulin-1 | Novus Biologicals | NBP1-84725 | IF: 1:100,<br>WB: 1:500 | 16487 |
| ECM1 | Proteintech Group Inc | 11521-1-AP | IF: 1:50,<br>WB: 1:500 | 59326 |
| Versican GAGβ | Sigma-Aldrich | AB1033 | IF: 1:50 | 3237533 |
| PCNA | Abcam | ab92552 | IF: 1:200 | GR3396553-19 |
| CD68 | Bio-Rad | MCA1957GA | IHC: 1:200 | 1708 |
| vWF | Dako | A0082 | IHC: 1:1000 | 111(101) |

**Supplemental Table 5.** Oligonucleotides for RT-PCR

| Gene Symbol | Accession No. | Forward (5′-3′)<br>Reverse (5′-3′) | Size (bp) |
|---|---|---|---|
| *Fbln1C* | NM_001347088.1 | AGAACTATCGCCGCTCCGCA<br>CCACCGCTGGCACTTGGATG | 135 |
| *Fbln1D* | EU543224.1 | TGAATGCCCCGAGAACTATC<br>CTCAGGACGGGTGAACTCTC | 178 |
| *Ecm1* | NM_007899.5 | GAGACCCTCAATGTGCTGGA<br>ATTGCATCCTCCCACACAAG | 101 |

**Supplemental Materials and Methods**

*Genotyping of EP4 reporter mice (Ptger4-IRES-nlsLacZ)*

DNA was genotyped by polymerase chain reaction (PCR) using a forward primer 5′-TGTTGGTGGATGAGGTTAGTGG-3′ (common primer), a wild-type reverse primer 5′-TCGCCTGTTCTCTAGTGGGA-3′, and a mutant-specific reverse primer 5′-CCTAGGAATGCTCGTCAAGAAGA-3′. The wild-type *Ptger4* allele was identified as a 212-bp amplicon, and the mutant *Ptger4* allele was identified as a 325-bp amplicon. Cycling parameters for PCR were 35 cycles of 94 °C for 30 s, 58 °C for 30 s, and 72 °C for 30s.

*Genotyping of Fbln1 floxed mice and Fbln1 exon 3 deleted mice*

DNA was genotyped by PCR using a forward primer 5′-CCTTGCTCCCACCCCGGACATC-3′ (common primer), a wild-type reverse primer 5′-CCAGAGAACCCCTGCTCTGCC-3′, and a mutant-specific reverse primer 5′-CAGAGCCTCAGAAGAACTGATGGCGAGC-3′. The wild-type *Fbln1* allele was identified as a 528-bp amplicon, and the *Fbln1* floxed allele was identified as a 628-bp amplicon. Cycling parameters for PCR were 29 cycles of 95 °C for 30 s, 55 °C for 30 s, and 72 °C for 1 min.

To genotype mice following excision of *Fbln1* exon 3 by Cre recombinase, DNA was amplified by PCR using the primers 5′-TGGATGAGAGCCATATTTGACATCCTTCCG-3′ and 5′-GAGGCCCTAGTGTGGGTCTGGTGTGAGC-3′. The wild-type *Fbln1* allele was identified as a 1026-bp amplicon, and the *Fbln1* allele missing exon 3 was identified as an approximately 300-bp amplicon. Cycling parameters for PCR were 29 cycles of 95 °C for 30 s, 55 °C for 30 s, and 72 °C for 1 min 30 s.

*Oral administration of an EP4 antagonist in mice*

CJ-42794 (0.1 mg/kg in 0.5% methylcellulose) was administered orally to C57BL/6N mice (male,12-14 weeks old) at a dosage of 0.5 mL/100 g of body weight, twice daily (equivalent to 0.2 mg/kg/day) for 4 weeks after vascular injury. CJ-42794 was prepared immediately before administration. Mice in the control group received the same dosage of 0.5% methylcellulose (Wako, Osaka, Japan).

*Tissue staining and evaluation of IH*

Paraffin-embedded blocks containing femoral artery tissues were sectioned to a thickness of 4 µm and placed on MAS-coated glass slides (Matsunami, Osaka, Japan). To elucidate the structural characteristics of the vessel wall, the sections were subjected to Elastica van Gieson (EVG) staining (Muto Pure Chemicals, Tokyo, Japan), following the manufacturer’s instructions. The evaluation of intimal hyperplasia was based on EVG-stained images, in which were analyzed with Image J software (version

1.53) to determine neointima area, medial layer area, neointima-to-medial layer ratio, medial layer thickness, length of the external elastic laminae, and internal lumen area.

*Immunofluorescent staining*

Immunofluorescent staining was performed following previously described protocols.[1,2] Co-staining of fibulin-1, ECM1, or versican GAGβ domain was conducted with Hoechst. Co-staining of PCNA was also conducted with Hoechst. Specifically, chondroitinase ABC treatment (Sigma-Aldrich) was carried out for 1 h at 37 °C exclusively for versican GAGβ. To activate antigenicity, fibulin-1 underwent incubation in a 10 mM citrate buffer (pH 6.0) for 10 min at 700 W in a microwave oven, followed by cooling to room temperature. ECM-1 and PCNA were incubated in a 1 mM EDTA solution (pH 9.0) at 95 °C for 30 min. Subsequently, the tissue samples were incubated in a 0.1% Triton X reagent (Sigma-Aldrich) for 10 min to enhance cell membrane permeability. Each primary antibody was added and kept at 4 °C for two consecutive overnight periods. The concentration of each antibody is provided in Supplemental Table 4. After incubation, the tissues were stained with Alexa Fluor 546 or 647 anti-rabbit IgG (dilution: 1:250) for 1 h. DNA was stained using Hoechst 33258 solution (dilution: 1:2000). The resulting images were acquired using an LSM 710 Confocal Microscope (Zeiss, Oberkochen, Germany).

*Immunohistochemistry*

αSMA, von Willebrand factor, or CD68 were co-stained with Nuclear Fast Red staining using XGal-stained specimens. Specimens were deparaffinized, rehydrated, and incubated. The tissue sections were then incubated in 0.3% $H_2O_2$ solution (Wako) for 30 min to inactivate endogenous peroxidase. To activate antigenicity of CD68 and αSMA, specimens were incubated in 10 mM citrate buffer (pH 6.0) for 10 min at 700 W in a microwave oven and cooled to room temperature. To activate antigenicity of von Willebrand factor, specimens were incubated in 20 $\mu$g/ml protein kinase K for 10 min at 37 °C. Specimens were then incubated with 1.5% serum (Vectastain Elite ABC IgG Kit: Vector Laboratories Burlingame, CA, USA) at room temperature for 30 min to block nonspecific reactions. Primary antibodies were added to the specimens and incubated at 4 °C overnight. Specimens were incubated with biotinylated anti-mouse/rabbit IgG (Vectastain Elite ABC IgG Kit: Vector) for 30 min and then avidin/biotinylated enzyme (Vectastain Elite ABC IgG Kit: Vector) for 30 min at room temperature. Finally, color development was performed using DAB solution (Peroxidase Stain DAB kit: Nacalai Tesque). All processes were washed 3 times with 0.1 M PBS for 5 min; then specimens were dehydrated, permeabilized, and sealed.

*Primary Culture of VSMCs*
Primary cultures of VSMCs isolated from *Ptger4*-Tg, non-Tg, *Fbln1*$^{fl/fl}$;*SM22-Cre*, and *Fbln1*$^{fl/fl}$ mice were obtained using an explant method as previously described.[3] Briefly, after euthanizing the mice via cervical dislocation following deep isoflurane inhalation (3% at a flow rate of 1 L/min), the descending aortas were collected, and the surrounding fat and connective tissue were removed. The aortas were then digested using a collagenase enzyme mixture consisting of 1.5 mg/mL collagenase-dispase (Roche, Basel, Switzerland), 2.0 mg/mL BSA fraction V (Sigma-Aldrich), 1 mg/mL trypsin inhibitor type I-S (Sigma-Aldrich), and 1 mg/mL collagenase II (Sigma-Aldrich) dissolved in HBSS (Nacalai Tesque) at 37 °C for 5 min to isolate the smooth muscle layers. The smooth muscle layers were plated onto poly-L-lysine (Wako)-coated dishes. These cells were cultured in Dulbecco's modified essential medium (DMEM, Nacalai Tesque) supplemented with 20% fetal bovine serum (FBS, Sigma-Aldrich) and penicillin-streptomycin (Wako) under 5% $CO_2$ at 37 °C.

*Quantitative Reverse Transcription-PCR*
Cells were plated in 12-well plates at a density of 7 × $10^4$ cells/well for RNA extraction. Before stimulation, the cells were serum-starved in DMEM without FBS for 24 h. To inhibit endogenous $PGE_2$ production by VSMCs, indomethacin was administered at 100 μmol/L for 1 h before stimulation. Cells were then stimulated with $PGE_2$ (1 μmol/L), $PGE_2$ (1 μmol/L) + ONO-AE3-208(1 μmol/L), or ONO-AE1-437(1 μmol/L) for 24 h.

Reverse transcription was performed using the PrimeScript RT reagent kit (TaKaRa Bio, Shiga, Japan), and quantitative reverse transcription-PCR (RT-PCR) was carried out using TB Green FAST qPCR mix (TaKaRa Bio). The expression level of each gene was calculated relative to that of 18S ribosomal RNA using the ΔΔCT method. The primer sequences used are provided in Supplemental Table 5. Each PCR cycle consisted of denaturation at 95 °C for 5 seconds and annealing at 60 °C for 10 seconds. A total of 40 cycles were performed.

*Western Blotting*
Cells were plated on 6-well plates at a density of 1.5 × $10^5$ cells/well for protein extraction from the cell supernatant. The western blotting methods were as previously described.[1] For the detection of fibulin-1 and ECM1 proteins, *Ptger4*-Tg VSMCs was incubated with ONO-AE1-329 (1 μmol/L) for 72 h in DMEM without FBS. Before stimulation, indomethacin was administered at 100 μmol/L for 1 h. Supernatants (2 mL) were concentrated to a volume ≤50 μL using a 10-K Amicon Ultra-2 mL Centrifugal Filter (Millipore, Burlington, MA, USA). Proteins were separated by SDS-PAGE and transferred to a polyvinylidene difluoride membrane via electroblotting. The membranes were blocked in TBS-0.1% Tween containing 5% skim milk and subsequently incubated

with anti-fibulin-1 antibody (dilution: 1:500) or anti-ECM1 antibody (dilution: 1:500) at 4 °C overnight. The membranes were then incubated with peroxidase-linked secondary antibodies (dilution: 1:5000; Cell Signaling Technology, Danvers, MA, USA) for 1 h. Chemiluminescence detection was performed using the ECL reagent (ThermoFisher Scientific). Signals were visualized using a LuminoGraph II (ATTO, Tokyo, Japan), and the band intensities were quantified using ATTO CS Analyzer 4 software (ATTO).

*Expression and Purification of Recombinant ECM1 Protein*

The full-length human ECM1 recombinant protein was obtained as described previously.[4]

*Proliferation Assay*

Cell proliferation was assessed using the cell counting kit-8 (Dojindo Laboratories, Kumamoto, Japan) as per the manufacturer's instructions. Non-Tg VSMCs were plated on 96-well plates at a density of $5 \times 10^3$ cells/well and then starved overnight in DMEM containing 0.5% FBS. Subsequently, fibulin-1C (1 μg/mL) or fibulin-1D recombinant proteins (1 μg/mL) were used in Figure 4H. Full-length or deletion mutant of fibulin-1C and ECM1 recombinant proteins with C-terminal His-tag were used for cell proliferation assay. The concentrations of the mutant proteins were calculated based on each molecular weight estimated by SDS-PAGE (Supplemental Figure 2). The concentrations of each recombinant proteins are as follows: (fibulin-1C, 1 μg/mL; ΔEG2-3, 0.9 μg/mL; ΔEG4-5, 0.9 μg/mL; ΔEG2-5, 0.72 μg/mL; ΔEG6-7, 0.9 μg/mL; ΔEG2-7, 0.63 μg/mL; ΔEG2-8, 0.54 μg/mL; ΔEG8-CT, 0.72 μg/mL; ECM1, 1 μg/mL). Cell proliferation was measured 48 h after adding recombinant proteins using the GloMax Explorer Multimode Microplate Reader (GM3500; Promega, Madison, WI, USA).

*Fbln1*$^{fl/fl}$ and *Fbln1*$^{fl/fl}$;*SM22-Cre* VSMCs were plated in 96-well plates at a density of $5 \times 10^3$ cells/well and then starved overnight in DMEM containing 0.5% FBS. Cells were serum-starved in DMEM with 0.5% FBS for 24 h and subsequently stimulated with ONO-AE1-437 (1 μmol/L). Indomethacin was administered at 100 μmol/L for 1 h before stimulation. The cells were measured 48 h later using the GM3500 (Figure 4G).
